# Restricted X chromosome introgression and support for Haldane’s rule in hybridizing damselflies

**DOI:** 10.1101/2021.11.28.470238

**Authors:** Janne Swaegers, Rosa Ana Sánchez-Guillén, Pallavi Chauhan, Maren Wellenreuther, Bengt Hansson

## Abstract

Contemporary hybrid zones act as natural laboratories for the investigation of species boundaries and allow to shed light on the little understood roles of sex chromosomes in species divergence. Sex chromosomes are considered to function as a hotspot of genetic divergence between species; indicated by less genomic introgression compared to autosomes during hybridisation. Moreover, they are thought to contribute to Haldane’s rule which states that hybrids of the heterogametic sex are more likely to be inviable or sterile. To test these hypotheses, we used contemporary hybrid zones of *Ischnura elegans*, a damselfly species that has been expanding its range into the northern and western regions of Spain, leading to chronic hybridization with its sister species *Ischnura graellsii*. We analysed genome-wide SNPs in the Spanish *I. elegans* and *I. graellsii* hybrid zone and found (i) that the X chromosome shows less genomic introgression compared to autosomes and (ii) that males are underrepresented among admixed individuals as predicted by Haldane’s rule. This is the first study in Odonata that suggests a role of the X chromosome in reproductive isolation.

Moreover, our data adds to the few studies on species with X0 sex determination system and contradicts the hypothesis that the absence of a Y chromosome causes exceptions to Haldane’s rule.

## INTRODUCTION

Since Darwin’s theory of evolution (Darwin 1859) it has become clear that speciation – the evolution of reproductive barriers between populations – is complex and continuous. It is already well established that due to independent assortment and recombination, genome regions have unique evolutionary histories. For example, alleles that are neutral or (generally) adaptive are expected to cross species boundaries, while alleles under divergent selection or associated with reproductive isolation do not (Ravinet et al. 2017). Species boundaries can therefore be expected to be ‘semipermeable’. The heterogeneity of genomic divergence is expected to be the result of the interplay between natural and sexual selection as well as gene flow, demography and recombination. However, characterizing the genomic architecture of barriers to gene exchange remains a key challenge in studies of speciation (Payseur and Rieseberg 2016; Fraïsse and Sachdeva 2020), especially in non-model species (Fraïsse and Sachdeva 2020).

Contemporary hybrid zones – regions where species hybridize and introgress – offer fascinating opportunities to study speciation (Gompert et al. 2017). First, hybrid zones act as natural laboratories for the investigation of species boundaries and more generally the origin of species (Harrison and Larson 2016). It is within these hybrid zones that divergent loci associated with reproductive isolation can be detected. This is in contrast to the comparison of allopatric (non-overlapping) parental species, where divergent loci can reflect different selection pressures and/or random effects operating after speciation has been completed (Nosil and Schluter 2011; Feder et al. 2013). Second, hybrid zones allow to shed light on the little understood role of sex chromosomes in facilitating species divergence. Indeed, loci that are showing divergence between species are expected to be enriched on sex chromosomes (the “large X-effect”) as recessive loci that increase fitness in the heterogametic sex (males in XY systems) would accumulate faster on the X chromosome because of immediate exposure to selection (Meisel and Connallon 2013). Other processes such as recombination rate, mutation rate and effective population size differences between X chromosomes and autosomes, may add to this pattern (Meisel and Connallon 2013; Charlesworth et al. 2018). The X chromosome is therefore considered to function as a hotspot of genetic divergence between species; indicated by less genomic introgression compared to autosomes during hybridization. Consequently, due to hybrid incompatibilities on sex chromosomes it can also be expected that the hybridised heterogametic sex suffers from a fitness reduction compared to the homogametic sex (“Haldane’s rule”; Haldane 1922).

Although vast evidence has been found for these processes, this evidence comes from a limited number of lineages (with a majority of studies in birds, Lepidoptera and Diptera (Presgraves 2018)), and sex determination systems (mainly XY and ZW systems (Presgraves 2018)). To expand our knowledge on the role of sex chromosomes in speciation, more comprehensive knowledge is needed in a wider range of taxa and other sex determination systems, such as the X0 and Z0 systems (Fraïsse and Sachdeva 2020).

We therefore sought to clarify the role of the X chromosome in the origin of reproductive barriers in an insect order with an X0 sex determination system. More specifically, we focus on the recently established hybrid zone in Spain between the damselfly sister species pair *Ischnura elegans* and *I. graellsii* (Sánchez-Guillén et al. 2012a). X-linked genes and their properties have recently been identified in *I. elegans* (Chauhan et al., 2021), yet it has not been investigated whether “speciation genes” can be more often found on the X chromosome in this insect order. Hybridisation between the two species is the consequence of the recent anthropogenic-driven range expansion of *I. elegans* into the northern and western regions of Spain (Sánchez-Guillén et al. 2011). Both species have been studied in exceptional detail for the last 20 years, providing access to a wealth of ecological and natural history data. Admixture analyses in the hybrid zone have revealed that the majority of *I. elegans* show levels of introgression similar to those expected for *I. elegans* backcrosses, and in a few cases F_1_ hybrids (first generation hybrids) (Sánchez-Guillén *et al.* 2011). *Ischnura* damselfly females have one pair of X chromosomes (XX), whereas males have a single X chromosome (and no Y chromosome). Thus, females have a diploid sex chromosome karyotype (XX) whereas males are hemizygous for X (X0). To our knowledge, so far only two other studies investigated introgression patterns between autosomes and the X chromosome in species with an X0 sex determination system (both in the insect order Orthoptera; Maroja et al., 2015; Moran et al., 2018). Interestingly, the absence of a Y chromosome might relax several mechanisms that contribute to Haldane’s rule, such as incompatibilities between Y-linked and autosomal genes (Sweigart 2010; Campbell et al. 2012) and meiotic drive (Patten 2018). By studying species with an X0 sex determination system we can explore whether these mechanisms are necessary for Haldane’s rule to apply. The few existing case studies of X0 sex determination systems show incidentally rare exceptions to Haldane’s rule (Moran et al. 2017a).

Today high-throughput sequencing technology provides unprecedented opportunities to study genomic evolutionary histories at hybrid zones (Harrison and Larson 2016) allowing exciting approaches to disentangle evolutionary processes across the speciation continuum (Gompert et al. 2017). Here we analyse genome-wide distributed single nucleotide polymorphisms (SNPs) in the Spanish *I. elegans* and *I. graellsii* hybrid zone to test whether the X chromosome shows less genomic introgression compared to autosomes and whether X0 males are underrepresented among hybrids and backcrosses as predicted by Haldane’s rule during hybridization caused by range expansion.

## METHODS

### Sampling strategy

We sampled individuals from fifteen localities in the hybrid zone along with five localities of allopatric *I. elegans* and three localities of allopatric *I. graellsii* (Fig. 1A; for details see Table S1 in Supplemental Information). Additionally, three closely related species from the *elegans-clade* (*I. fountaineae, I. genei* and *I. saharensis*) were also sampled (Table S1 in Supplemental Information).

**Figure 1.**
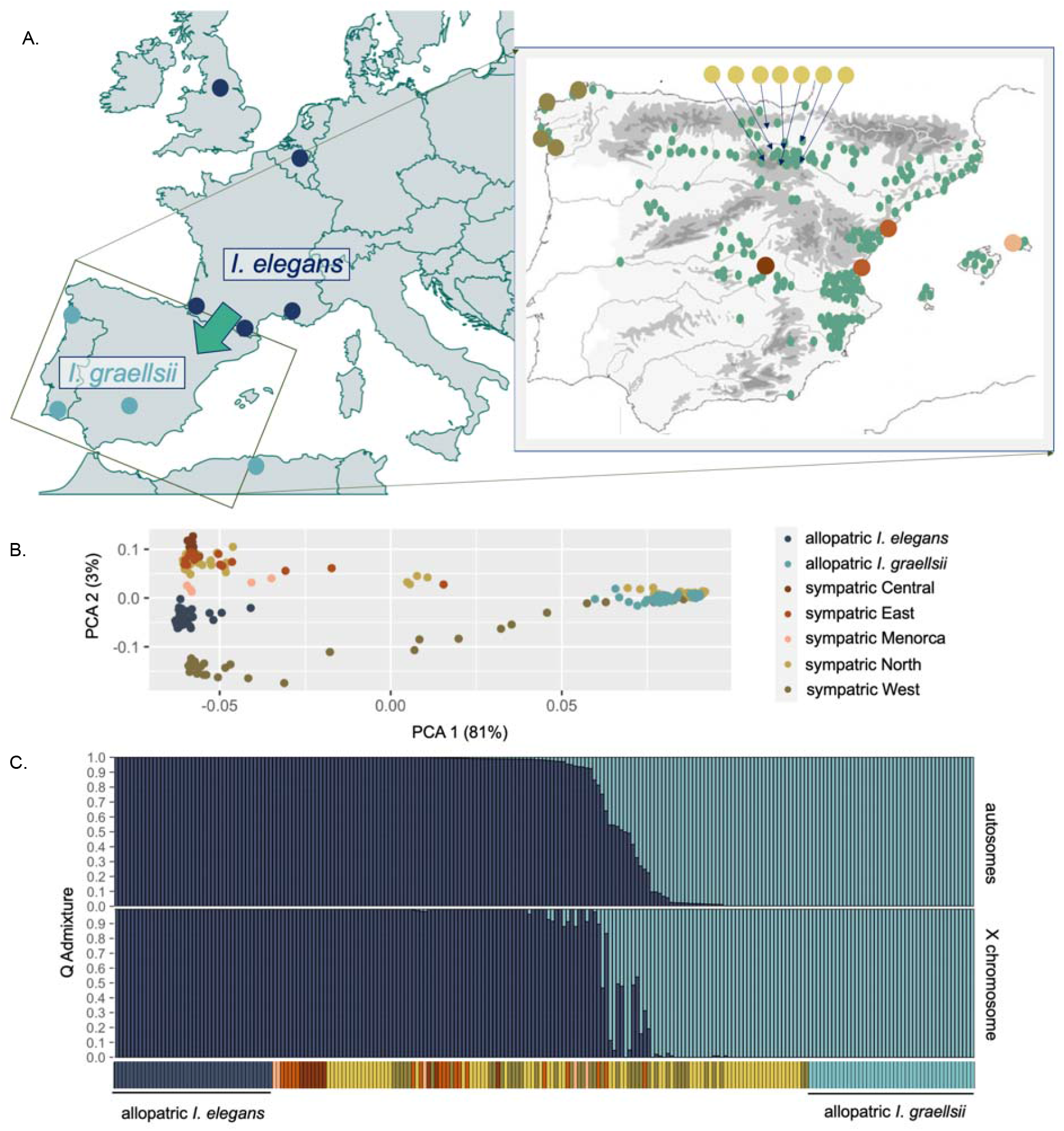
(A) Maps showing the allopatric (left) and sympatric populations (right) of *I. elegans* and *I. graellsii* that were studied. Green areas on the rightmost map indicate where *I. elegans* has expanded its range into Spain. (B) The first two axes of a principal component analysis (PCA) of all allopatric and sympatric individuals. The colors match the sample locations on the map. (C) Individual admixture proportions (Q values) based on autosomal and X-linked SNPs, respectively. Samples have been ordered based on the Q values from autosomal SNPs.

### Library construction, RAD-seq analysis and filtering

Genomic DNA from the head and thorax of 269 individuals (260 samples of *I. elegans* and *I. graellsii* and nine samples of closely related *Ischnura* species that were used as outgroup samples in part of subsequent analyses, Table S1 in Supplementary Information) was extracted with the DNeasy Blood & Tissue Kit (Qiagen). Extracted genomic DNA was quantified using Nanodrop and Qubit and DNA degradation was visually expected through 1% agarose gel electrophoresis. In total, eight single-digest Restriction site–Associated DNA (RAD) libraries were constructed following the protocol described in Etter et al. (2011) and modified in Dudaniec et al. (2018). Per library, forty unique barcodes were used to label the samples (sourced from Metabion). Five of these libraries (containing 206 samples) were paired-end sequenced (2*100 bp) on separate lanes of an Illumina HiSeq 2500 at SNP&SEQ Technology Platform at Uppsala University, whereas the remaining three libraries (containing 61 samples) were paired-end sequenced (2*125 bp) on three lanes of an Illumina HiSeq 2500 at BGI (Hong Kong).

We used the bioinformatic pipelines in STACKS v2.2 (Catchen et al. 2011, 2013) to process the sequences. Process_radtags was used to demultiplex the raw reads, and clone_filter to identify and discard PCR clones using default parameters. Next, sequence reads were aligned to the *I. elegans* draft genome assembly (Chauhan et al. 2021) using BOWTIE2 v.2.3 (mismatch allowance per seed alignment of 1, maximum mismatch penalty of 6 and minimum of 2, maximum fragment length of 1000 bp and minimum of 100 bp, Langmead & Salzberg, 2012). The aligned samples were processed with the ref_map pipeline to detect SNPs using default parameters (different runs were performed when including and excluding outgroup samples).

We discarded 35 samples that had a mean depth < 20x and also two *I. graellsii* samples from the population Seyhouse (Algeria) as exploratory analyses of population structure revealed possible hybridization in those samples with a third *Ischnura* species (Sánchez-Guillén et al., in review). We generated three different SNP sets for subsequent analyses using ‘populations’ in STACKS: a first set including all SNPs detected among allopatric samples of *I. elegans* and *I. graellsii*; a second set with only diagnostic SNPs between the allopatric samples of *I. elegans* and *I. graellsii* (i.e. loci that are differentially fixed between these two groups), and a third set when also outgroup samples were included. For all three SNP sets, only SNPs with a minor allele frequency of > 0.05 and an observed heterozygosity of < 0.7 were retained. Moreover, loci had to occur in 80% of the individuals in a population. For the two non-diagnostic SNP sets, the locus had to occur in 80% of the individuals in a population and in 20 of the 25 (or 28 for the SNP set with outgroup samples included) populations to be included in the final SNP set. The SNP sets that did not include the outgroup samples were subsequently filtered to include only one random SNP per RAD-tag to create data without closely linked loci (using the write_random_snp option in STACKS). These SNP sets are hereafter referred to as the ‘full SNP set’ and the ‘diagnostic SNP set’, respectively, while the SNP set with the outgroup samples is referred to as the ‘outgroup SNP set. For all three SNP sets, we differentiated between SNPs that were located on autosomes versus the X chromosome based on an *I. elegans* reference genome assembly (Chauhan et al. 2021).

Next, we genotypically classified individuals as male or female based on observed homozygosity (H_O_) at X-linked SNPs. As males are hemizygous, we expect an H_O_ = 1.0 at X-linked SNPs for males, yet in practice deviations are expected due to genotyping error. As females have two copies of the X chromosome, we expect lower H_O_ in females compared to males. Accordingly, using data of X-linked SNPs at the full SNP set, we found that the H_O_ values among all *I. elegans* and *I. graellsii* samples were bimodally distributed (Figure S1 in Supplemental Information). We selected a cut-off value at the valley of the bimodal H_O_ distribution (i.e. H_O_ = 0.96) to classify samples having H_O_ < 0.96 as females and samples having H_O_ > 0.96 as males. In this way, we genotypically classified the *I. elegans* and *I. graellsii* samples as 129 females and 94 males. As we used samples that were in many cases > 10 years old, phenotypically sexing of individuals was not always straightforward. Among the *I. elegans* and *graellsii* samples that had been phenotypically classified as females all 96 had H_O_ < 0.96 as expected, whereas 19 of 105 phenotypically classified as males had H_O_ < 0.96 (these were treated as females in the analyses). Among the samples that had not been phenotypically sexed, 14 were classified as females and eight as males based on H_O_. The outgroup samples were genotypically classified as eight females and one male using H_O_ at X-linked SNPs at the outgroup SNP set.

Finally, we filtered the X-linked SNPs further by retaining only those SNPs that were homozygous in all genotypically classified males. This was done for all three SNP sets, giving the final SNP sets: the full SNP set with 7,352 SNPs of which 390 are X-linked, the diagnostic SNP set with 1,931 SNPs of which 111 are X-linked, and the outgroup SNP set with 64,452 SNPs of which 4,603 are X-linked. When analyses are performed on only autosomal SNPs or only X-linked SNPs, we referred to these SNP sets as, e.g., the X-linked full SNP set or the autosomal diagnostic SNP data set.

### Population structure analysis

To discern population structure among the samples, we performed Principal Component Analysis (PCA) using the PCA function in PLINK v1.9 (Purcell et al. 2007). For this analysis, we used autosomal SNPs from the full SNP set).

### Individual ancestry coefficients

We compared the ancestry of individuals to allopatric *I. elegans* and *I. graellsii* between autosomes and the X chromosome by calculating individual ancestry coefficients (*Q-* values) using both the autosomal and X-linked diagnostic SNP set in ADMIXTURE v1.3.0 (Alexander and Lange 2011). ADMIXTURE was run using the supervised learning mode with the allopatric *I. elegans* and *I. graellsii* individuals as reference samples meaning 100% ancestry is assumed for the respective species. For the X-linked diagnostic SNP set, hemizygosity was accounted for by setting the haploid flag for all males.

### Introgression analysis

We used two different approaches to infer whether introgression patterns are different between autosomes versus the X chromosome. First, we employed a Bayesian Genomic Clines (BGC) analysis of Gompert & Buerkle (2011, 2012), which makes use of Markov chain Monte Carlo to estimate genomic cline parameters within a Bayesian genomic cline model. The per locus probability of being inherited from a given parental population (⍰) is calculated, which is then compared to the genome-wide average probability, i.e. the hybrid index. Two parameters, α and β, summarise this probability and hence the pattern of introgression between the parental populations that are nearly fixed for the focal markers. For this analysis we used the autosomal and X-linked diagnostic SNP set. In our case, the parameter α measures the directional movement of alleles from *I. graellsii* into *I. elegans* (α > 0) or movement from *I. elegans* into *I. graellsii* (α < 0), while the β parameter, measures the strength of the barrier to gene flow between the two species. Higher positive values of the β parameter describe steeper clines and a greater strength of the gene flow barrier. We ran 5 independent chains in BGC using the genotype certainty model, each for 50,000 steps with a burn-in of 25,000 and thinning samples by 20. We combined the output for both α and β using ClineHelpR (available at https://github.com/btmartin721/ClineHelpR). To test whether X-linked SNPs displayed higher β values than the autosomes, we generated 10,000 permuted datasets by sampling without replacement from the autosomal β value distribution. For each dataset we sampled 111 times, i.e. the number of X-linked diagnostic SNPs, to generate equal sample sizes between autosomal and X-linked datasets. Subsequently, we compared the median of β values of the X-linked distribution to the median of each permuted autosomal dataset and considered a greater gene flow barrier on X-linked SNPs compared to autosomal SNPs if the X-linked observed median exceeded the median in > 95% of the permuted datasets (Baiz et al. 2020).

Second, we made use of ABBA-BABA statistics which are based on the relative frequency of shared alleles between three focal groups, along one outgroup to determine which allele is ancestral. In our case, we compare (i) the frequency of shared alleles between sympatric *I. elegans* and allopatric *I. graellsii* (‘ABBA’) compared to shared alleles between allopatric *I. elegans* and allopatric *I. graellsii* (‘BABA’), and (ii) the frequency of shared alleles between sympatric *I. graellsii* and allopatric *I. elegans* (‘ABBA’) compared to shared alleles between allopatric *I. graellsii* and allopatric *I. elegans* (‘BABA’) (see Figure 3). If introgression occurs in sympatry, higher frequencies of ABBA than of BABA are expected. Patterson’s D is the original test statistic used to measure this but is now often used in parallel with related test statistics *f*_d_ and *f*_dM_ that are less biased when, for example, used in sliding windows frameworks (Malinsky et al. 2021). We here report the results using *f*_dM,_ yet similar results were found with test statistics D and *f*_d_ (results not given). For this analysis we used the outgroup SNP set. We ran Dsuite (Malinsky et al. 2021) to measure these test statistics along the genome using a sliding window approach. More specifically, we ran the function Dinvestigate with a window size of 50 informative SNPs and a step of 5 SNPs. As outgroup we used 3 samples each from congeneric species *I. genei, I. fountaineae* and *I. saharensis.* We calculated the introgression parameters both for introgression from allopatric *I. graellsii* into sympatric *I. elegans* and from allopatric *I. elegans* into sympatric *I. graellsii*. In Figure 3 is depicted which samples we used as ‘P1’, ‘P2’ and ‘P3’ for both analyses (‘P4’ is the outgroup). As we wanted to include sympatric individuals that can be considered to be genomically *I. elegans* or *I. graellsii*, respectively, in this analysis, but did not know how incorporating individuals of more recent hybrid ancestry will affect the results, we used different autosomal Q admixture cut-off values to decide which sympatric individuals can be considered to be either genomically *I. elegans* or *I. graellsii.* (i) Q = 0 for sympatric *I. elegans* and Q = 1 for sympatric *I. graellsii*, (ii) Q < 0.1 for sympatric *I. elegans* and Q > 0.9 for sympatric *I. graellsii*, (iii) Q < 0.25 for sympatric *I. elegans* and Q > 0.75 for sympatric *I. graellsii*. We ran one analysis for each of these chosen cut-off values per species (six analyses in total).

**Figure 2.**
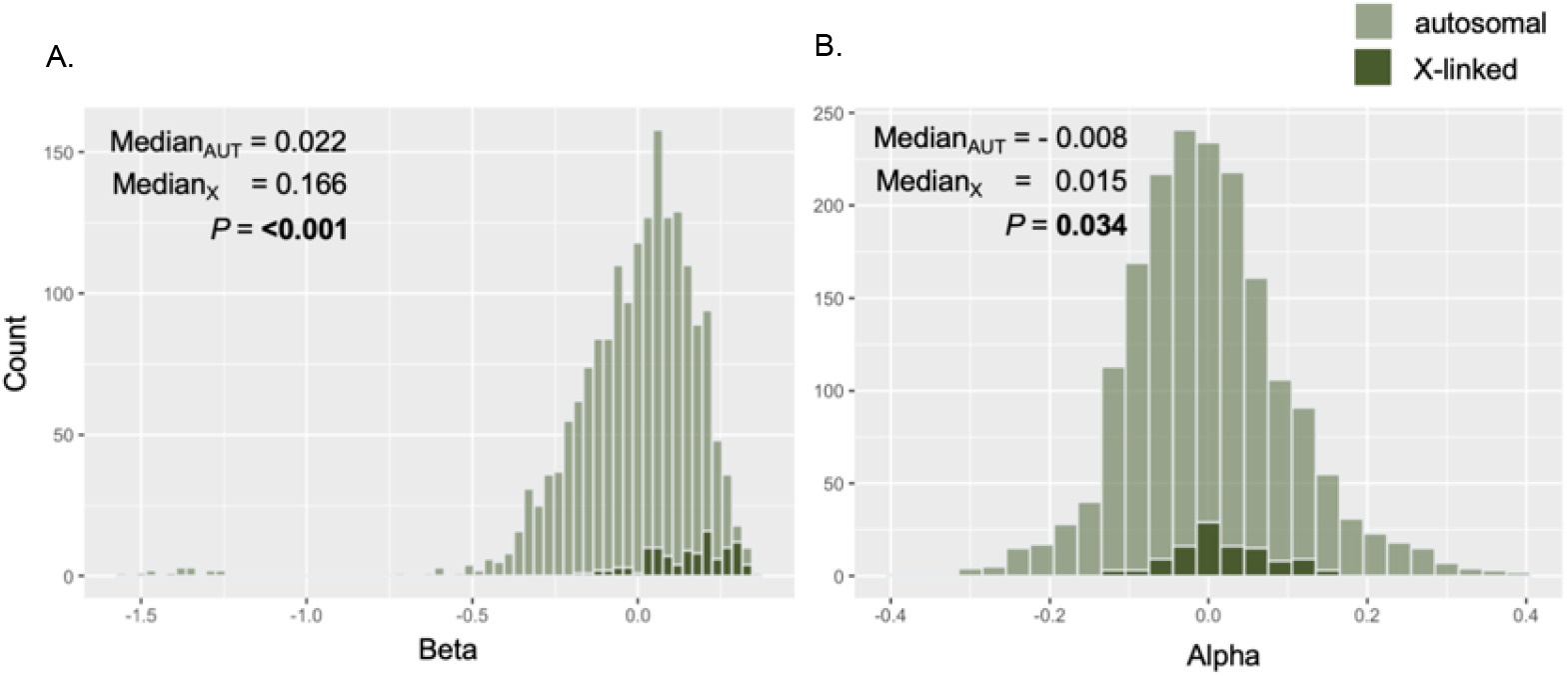
Results of BGC analysis of females in sympatric populations of *I. elegans* and *I. graellsii*. Shown are the beta (A) and alpha (B) distributions. The medians of the distributions measured autosomal and X-linked SNPs, respectively, are given in each panel, as well as the *P-*value from a permutation test comparing these medians.

**Figure 3.**
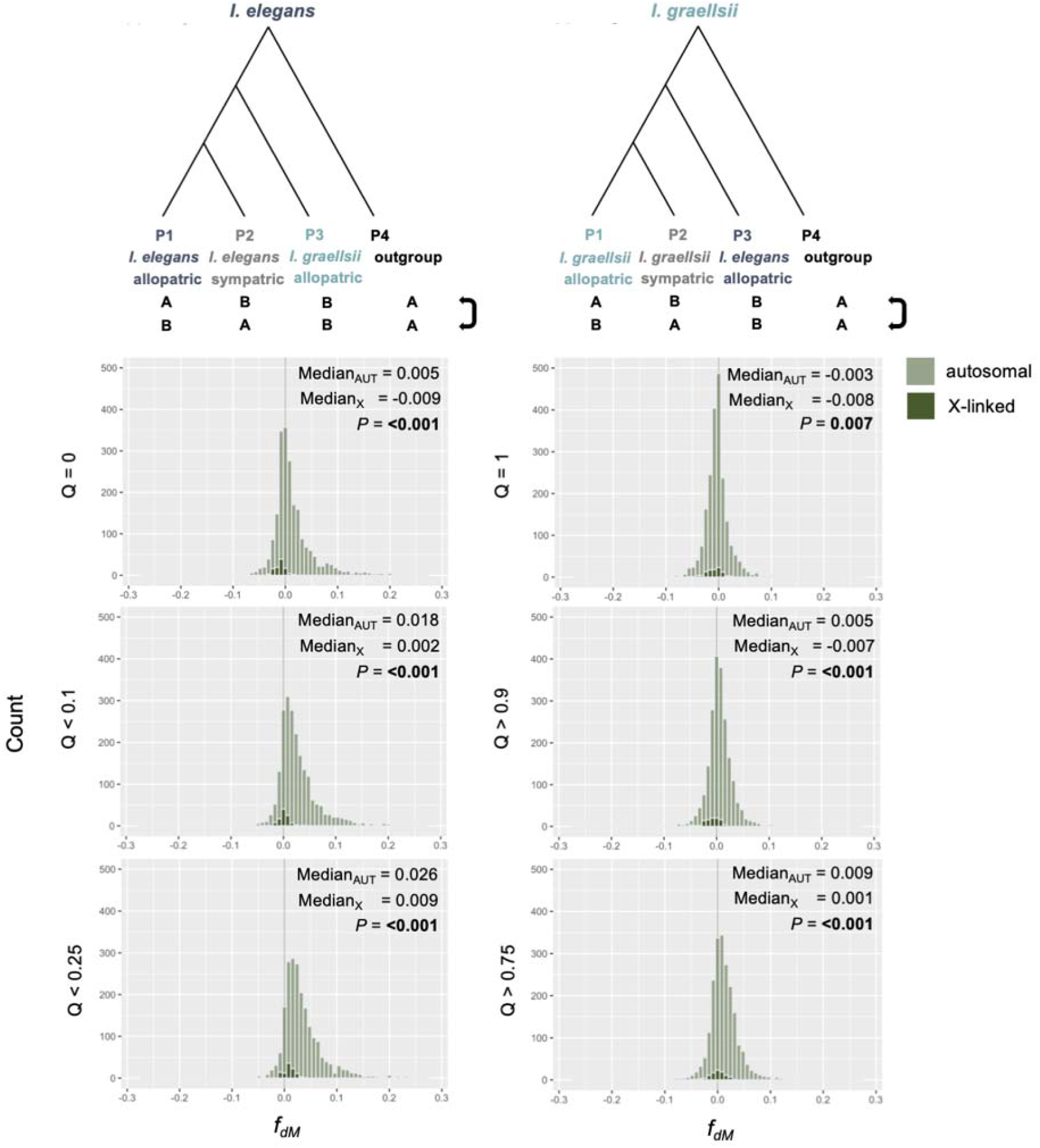
Results of ABBA-BABA analysis of females in sympatric populations of *I. elegans* and *I. graellsii*. Shown are the *f*_dM_ distributions. The medians of the distributions measured with autosomal and X-linked SNPs, respectively, are included in each panel, as well as the *P-* value from a permutation test to compare these medians. Left panels show results from sympatric *I. elegans*, and right panels sympatric *I. graellsii*.

Analogously to the BGC analysis, we generated permuted datasets from the distributions of test statistics of the autosomal windows and compared the medians of these to the median of the test statistics of the observed X-linked distribution. This was done for all six analyses with the given autosomal Q admixture cut-off value. Note that using the autosomal Q admixture cut-off value is a conservative approach to compare introgression levels between autosomal and X-linked windows. We considered there to be less introgression on the X chromosome compared to autosomes if the X-linked observed median was less than the median in > 95% of the permuted datasets.

As it is not possible to analyse males as hemizygous at the X chromosome in BGC and Dsuite, we ran these analyses using a subset of the data containing only the genotypically classified female individuals. However, we reran the analyses including both males and females (which did not change the results qualitatively; see below).

### Haldane’s rule

To test whether males were underrepresented among sympatric admixed individuals, we tested for associations between sex and proportion admixture for three different autosomal and X-linked Q admixture cut-off values using the full SNP set. As only females were sampled in the Western sympatric region (‘sympatric West’, Fig. 1A) we excluded all individuals in this region from the analysis. The following cut-off values were used to differentiate between admixed and non-admixed individuals: (i) Q = 0 or 1 (non-admixed individuals) and 0 < Q < 1 (lowly to highly admixed individuals), (ii) 0.1 > Q > 0.9 (non- to lowly admixed individuals) and 0.1 < Q < 0.9 (moderately to highly admixed individuals), and (iii) 0.25 > Q > 0.75 (non- to moderately admixed individuals) and 0.25 < Q < 0.75 (highly admixed individuals). Fisher’s exact tests were used to test whether males and females differed in numbers of non-admixed and admixed individuals for each Q value cut-off based on autosomal and X-linked SNPs, respectively.

## RESULTS

### Genetic structure

A principal component analysis of all allopatric and sympatric *I. elegans* and *I. graellsii* individuals based on autosomal SNPs at the full SNP set clearly separated the allopatric populations at the first axis (PC1) which explained much of the variation (Figure 1B). In contrast, some of the sympatric populations in the hybrid zone spread out along PC1, and separated partly along the minor second axis, PC2 (Figure 1B). An admixture analysis confirmed these patterns by grouping individuals in allopatric populations in separate clusters, while some sympatric samples had intermediate admixture proportions (Q values; Figure 1C). Interestingly, more individuals had intermediate Q values using autosomal SNPs compared to when using X-linked SNPs. At X-linked SNPs, sympatric individuals were more often showing Q admixture values closer to the values of allopatric individuals (Figure 1C).

### Bayesian genomic clines

We tested the strength and direction of allele movements between species using the diagnostic SNP set in females. We found that β values were significantly higher at X-linked SNPs compared to autosomal SNPs (permutation test, *P* < 0.001; Figure 2A). Also, the α parameter was higher at X-linked compared to autosomal SNPs (*P* = 0.034 Figure 2B). In other words, X-linked SNPs showed steeper clines (and hence a greater strength of the gene flow barrier) with alleles more likely to move from *I. graellsii* into *I. elegans* compared to the autosomal SNPs. Indeed 87% of the X-linked SNPs showed positive β values compared to 56% of the autosomal SNPs and 59% showed positive α values compared to 46% in the autosomes. Similar results were found in analysis that included both females and males (Table S3 in Supplementary Information).

### ABBA-BABA

Figure 3 shows the distributions of *f*_dM_ statistics between autosomal windows and windows located on the X chromosome. This statistic has the advantage of being symmetrically distributed around zero under the null hypothesis of no introgression and quantifies shared variation between P2 and P3 (positive values; ABBA) or between P1 and P3 (negative values; BABA) equally. For most Q admixture cut-off values used to include sympatric individuals (i.e., Q = 0 or 1; Q < 0.1 or > 0.9; Q < 0.25 or > 0.75), X-linked SNPs showed significantly less introgression (*f*_dM_ values distributed close to 0) between allopatric *I. graellsii* and sympatric *I. elegans* (*I. elegans* panel), and between allopatric *I. elegans* and sympatric *I. graellsii* (*I. graellsii* panel), than autosomal SNPs (*f*_dM_ biased towards positive values; permutation test, *P* ≤ 0.01 in all six analyses; Figure 3). Overall, the bias towards more introgression of autosomal than X-linked SNPs was more apparent for introgression into sympatric *I. elegans* (*I. elegans* panel). From Figure 3 it can also be concluded that overall introgression occurs more frequently from allopatric *I. graellsii* into sympatric *I. elegans* than from allopatric *I. elegans* into sympatric *I. graellsii* (Wilcoxon rank sum test, *P* < 0.001 in all three comparisons, Q = 0 vs Q = 1; Q < 0.1 vs. Q > 0.9; Q < 0.25 vs. Q > 0.75). These above results are for analysis with females only, but similar results were found in analyses including also males (Table S3 in Supplementary Information).

### Haldane’s rule

Admixed males were overall underrepresented within the hybrid zone, but the degree of underrepresentation differed for autosomal and X-linked diagnostic SNPs, and when different admixture cut-off values were used to categorize individuals as admixed or non-admixed (Figure 4). For autosomal SNPs, males were significantly underrepresented in the admixed category both when individuals with Q values between 0.1 and 0.9 (0.1 < Q < 0.9; Fisher’s exact test, *P* < 0.007), and between 0.25 and 0.75 (0.1 < Q < 0.9; *P* = 0.050), were categorized as admixed (Figure 4, upper panels). However, when Q values between 0 and 1 (0 < Q < 1) were used to categorize admixed individuals, males were not significantly underrepresented among admixed individuals (*P* = 1). For X-linked SNPs, males were significantly underrepresented among admixed individuals when individuals with Q values between 0 and 1 (0 < Q < 1; *P* = 0.009), and between 0.1 and 0.9 (0.1 < Q < 0.9; *P* < 0.001), were categorized as admixed (Figure 4, lower panels). For Q values between 0.25 and 0.75 (0.25 < Q < 0.75), males were not significantly underrepresented among the admixed individuals (*P* = 0.120), but it should be noted that the numbers of sampled highly admixed individuals was very low (Figure 4).

**Figure 4.**
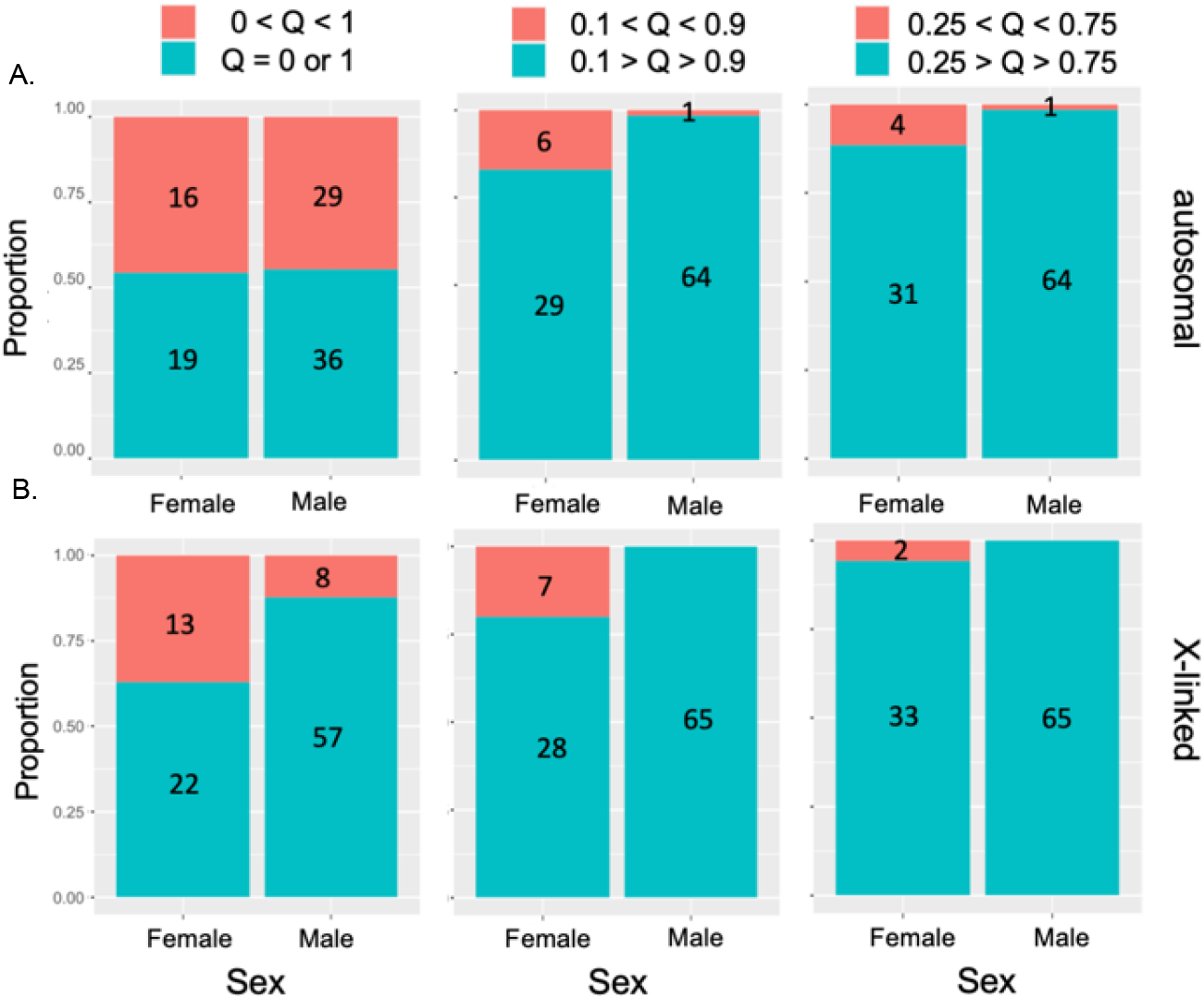
Proportion of admixed and non-admixed individuals in females and males, respectively, when using three different Q admixture cut-off values and either (A) autosomal and (B) X-linked SNPs. In all cases, males were underrepresented in the admixed (or highly admixed) category compared to females (Fisher’s exact test; A: *P* = 1, < 0.007, 0.050, respectively; B: *P* = 0.009, < 0.001, 0.120, respectively).

## DISCUSSION

In this study we analysed genome-wide distributed SNPs in the Spanish *I. elegans* and *I. graellsii* hybrid zone and found (i) that the X chromosome showed less genomic introgression compared to autosomes and (ii) that males are underrepresented among hybrids and backcrosses as predicted by Haldane’s rule.

### Introgression patterns

Through two different approaches and SNP sets (BGC using the ‘diagnostic SNP set’ with only *I. elegans* and *graellsii* samples, and ABBA-BABA using the ‘outgroup SNP set’ which also included outgroup samples) we detected lower introgression at the X chromosome compared to autosomes. Indeed, both methods measure introgression, yet BGC is a model-based approach while ABBA-BABA measures statistics proportional to the effective migration rate (Martin and Jiggins 2017). The similar results should be considered as complementary evidence for greater divergence at the X chromosome compared to autosomes between the two species in the hybrid zone. This is the first study in Odonata that suggests a role of the X chromosome in reproductive isolation. Although at this point direct evidence is lacking, our results suggest that a large X-effect may have contributed to an accumulation of reproductive barrier genes on the X chromosome. Only two other studies investigated introgression patterns between autosomes and the X chromosome in hybrid zones of species with an X0 sex determination system (both in insect order Orthoptera, Maroja et al., 2015; Peter A. Moran et al., 2018). Both these studies also detected that large X evolution has contributed to an accumulation of reproductive isolating genes on the X chromosome, as was detected in the current damselfly system.

Interestingly, both introgression analyses suggest that the direction of introgression is biased towards introgression from allopatric *I. graellsii* into sympatric *I. elegans*. This can be explained by two processes. First, from previous research in western Spain we know that there is asymmetry in the strength of the reproductive barriers between reciprocal crosses. Male *I. elegans* can more easily mate and produce hybrids with female *I. graellsii* and female hybrids, than the other way around (Sánchez-Guillén et al. 2012b, 2014). Curiously, this was also reflected here by the fact that the only sampled male F1 hybrid (autosomal SNP Q value: 0.5) had inherited its X chromosome from an *I. graellsii* mother. Overall weaker reproductive barrier in *I. elegans* would imply easier introgression into this species. Second, it could be expected that, in this case, alleles from *I. graellsii* rather than from *I. elegans* confer higher fitness in hybrid individuals (Gompert et al. 2017). This hypothesis is based on the rational that alleles from the native *I. graellsii* are expected to contribute more to local adaptation than those from *I. elegans* (which is relatively new to this region) (Wellenreuther et al. 2018). Note that even when reproductive barriers are strong between two species, adaptive introgression is possible (Gompert et al. 2017). Both mechanistic asymmetry as well as adaptive introgression could have acted simultaneously to the observed asymmetric introgression.

### Evidence for Haldane’s rule

When we compared the proportion of admixed versus non-admixed individuals between the sexes, we found fewer males than females among the admixed individuals. This pattern was pronounced at low levels of admixture of the X chromosome but not of the autosomes. Lower survival of males carrying hybrid and backcrossed X chromosomes is in accordance with the expectations from Haldane’s rule. Our data hence suggest that Haldane’s rule is valid in this insect order. An increased rate of mortality among hybrid and backcrossed males could be caused by the expression of recessive, deleterious alleles on the X chromosome in X0 male hybrids and homozygous females only, but not in heterozygous females (the latter which are more common). The observed pattern of stronger isolation between the two studied species at the X chromosome as compared to the autosomes further supports the presence of X-linked incompatibilities. This is one of the rare studies using a natural system in which the study species do not have a Y chromosome, and our results imply that neither incompatibilities between Y-linked and autosomal genes, nor meiotic drive, are necessary to cause the deleterious effects in male hybrids. Thus, our study does not support the suggestion that the absence of a Y chromosome constitute an exception to Haldane’s rule (Moran et al. 2017b).

Interestingly, the overall lower survival of males in the hybrid zone could impact sex-ratios and hence sexual conflict (Runemark et al. 2018). In the current species, sexual conflict over optimal mating rates is extensively studied (Sánchez-Guillén et al. 2017) and our results hence warrant further investigation on the effects of hybridization on sexual conflict.

### Conclusions

As predicted by theory, we here demonstrate that X-linked SNPs introgress less than autosomal SNPs in *I. elegans* and *I. graellsii* in the contemporary hybrid zone in Spain. Moreover, our data also suggest that Haldane’s rule is valid in Odonata and contradicts the hypothesis that the absence of a Y chromosome causes exceptions to Haldane’s rule. Thus, this is the first study in this insect order that suggests a role of the X chromosome in reproductive isolation. Future work is needed to establish if this also extends to other odonates and is thus a general rule. Expanding knowledge in the area of reproductive barriers, and mechanisms that fuel species melting, is urgently needed to predict biodiversity consequences under a scenario of climate induced range shifts that will increase the encounters of closely related species, and consequently the likelihood of introgressive hybridization (Sánchez-Guillén et al. 2016). Moreover, deciphering the relative contributions of X chromosomes and autosomes in keeping species together or not is shedding important fundamental insights into genome function and the evolutionary processes at play that contribute to speciation.

## ACKNOWLEDGEMENTS

We thank Adolfo Cordero Rivera, Iñaki Mezquita, Tomás Latasa, Mario García-París, Bernat Garrigós, Pere Luque, Xoaquín Baixeras, Francisco Cano, Jean Pierre Boudot, Jürgen Ott, Cedrick Vanappelghem, Philippe Lambret, and Phill Watts for their help with collecting the samples. Sequencing was performed by the SNP&SEQ Technology Platform at Uppsala Genome Center, which is part of National Genomics Infrastructure (NGI) Sweden, and Science for Life Laboratory (SciLifeLab) supported by the Swedish Research Council (and its Council for Research infrastructure, RFI) and the Knut and Alice Wallenberg Foundation. Bioinformatics analyses were performed on computational infrastructure provided by the Swedish National Infrastructure for Computing (SNIC) at Uppsala Multidisciplinary Center for Advanced Computational Science (UPPMAX).

## REFERENCES

Alexander, D. H., and K. Lange. 2011. Enhancements to the ADMIXTURE algorithm for individual ancestry estimation. BMC Bioinformatics 12:246.

Baiz, M. D., P. K. Tucker, J. L. Mueller, and L. Cortés-Ortiz. 2020. X-linked signature of reproductive isolation in humans is mirrored in a howler monkey hybrid zone. J. Hered. 111:419–428.

Campbell, P., J. M. Good, M. D. Dean, P. K. Tucker, and M. W. Nachman. 2012. The contribution of the Y chromosome to hybrid male sterility in house mice. Genetics 191:1271–1281.

Catchen, J., P. A. Hohenlohe, S. Bassham, A. Amores, and W. A. Cresko. 2013. Stacks: An analysis tool set for population genomics. Mol. Ecol. 22:3124–3140.

Catchen, J. M., A. Amores, P. Hohenlohe, W. Cresko, and J. H. Postlethwait. 2011. Stacks: Building and genotyping loci de novo from short-read sequences. G3 Genes, Genomes, Genet., doi: 10.1534/g3.111.000240.

Charles Darwin. 1859. On the origin of species. John Murray, London, UK.

Charlesworth, B., J. L. Campos, and B. C. Jackson. 2018. Faster-X evolution: theory and evidence from *Drosophila*. Mol. Ecol. 1–19.

Chauhan, P., J. Swaegers, R. A. Sánchez-Guillén, E. I. Svensson, M. Wellenreuther, and B. Hansson. 2021. Genome assembly, sex-biased gene expression and dosage compensation in the damselfly Ischnura elegans. Genomics 113:1828–1837.

Etter, P. D., S. Bassham, P. A. Hohenlohe, E. A. Johnson, and W. A. Cresko. 2011. SNP discovery and genotyping for evolutionary genetics using RAD sequencing. Methods Mol. Biol. 772:157–178.

Feder, J. L., S. M. Flaxman, S. P. Egan, A. A. Comeault, and P. Nosil. 2013. Geographic mode of speciation and genomic divergence. Annu. Rev. Ecol. Evol. Syst. 44:73–97. Annual Reviews.

Fraïsse, C., and H. Sachdeva. 2020. The rates of introgression and barriers to genetic exchange between hybridizing species: sex chromosomes vs. autosomes. Genetics, doi: 10.1101/2020.04.12.038042.

Gompert, Z., E. G. Mandeville, and C. A. Buerkle. 2017. Analysis of Population Genomic Data from Hybrid Zones. Annu. Rev. Ecol. Evol. Syst. 48:207–229.

Harrison, R. G., and E. L. Larson. 2016. Heterogeneous genome divergence, differential introgression, and the origin and structure of hybrid zones. Mol. Ecol. 25:2454–66.

Langmead, B., and S. L. Salzberg. 2012. Fast gapped-read alignment with Bowtie 2. Nat. Methods 9:357–9. Nature Publishing Group.

Malinsky, M., M. Matschiner, and H. Svardal. 2021. Dsuite - Fast D-statistics and related admixture evidence from VCF files. Mol. Ecol. Resour. 21:584–595.

Maroja, L. S., E. L. Larson, S. M. Bogdanowicz, and R. G. Harrison. 2015. Genes with restricted introgression in a Field Cricket (*Gryllus firmus/Gryllus pennsylvanicus*) hybrid zone are concentrated on the X chromosome and a single autosome. G3 Genes|Genomes|Genetics 5:2219–2227. Genetics Society of America.

Martin, S. H., and C. D. Jiggins. 2017. Interpreting the genomic landscape of introgression. Curr. Opin. Genet. Dev. 47:69–74. Elsevier Ltd.

Meisel, R. P., and T. Connallon. 2013. The faster-X effect: integrating theory and data. Trends Genet. 29:537–544.

Moran, P. A., S. Pascoal, T. Cezard, J. E. Risse, M. G. Ritchie, and N. W. Bailey. 2018. Opposing patterns of intraspecific and interspecific differentiation in sex chromosomes and autosomes. Mol. Ecol. 27:3905–3924.

Moran, P. A., M. G. Ritchie, and N. W. Bailey. 2017a. A rare exception to Haldane’s rule: Are X chromosomes key to hybrid incompatibilities? Heredity (Edinb). 118:554–562.

Moran, P. A., M. G. Ritchie, and N. W. Bailey. 2017b. A rare exception to Haldane ’ s rule: Are X chromosomes key to hybrid incompatibilities? 554–562.

Nosil, P., and D. Schluter. 2011. The genes underlying the process of speciation. Trends Ecol. Evol. 26:160–7.

Patten, M. M. 2018. Selfish X chromosomes and speciation. Mol. Ecol. 27:3772–3782.

Payseur, B. A., and L. H. Rieseberg. 2016. A genomic perspective on hybridization and speciation. Mol. Ecol. 25:2337–60.

Presgraves, D. C. 2018. Evaluating genomic signatures of “the large X-effect” during complex speciation. Mol. Ecol. 27:3822–3830.

Purcell, S., B. Neale, K. Todd-Brown, L. Thomas, M. A. R. Ferreira, D. Bender, J. Maller, P. Sklar, P. I. W. De Bakker, M. J. Daly, and P. C. Sham. 2007. PLINK: A tool set for whole-genome association and population-based linkage analyses. Am. J. Hum. Genet. 81:559–575.

Ravinet, M., R. Faria, R. K. Butlin, J. Galindo, N. Bierne, M. Rafajlović, M. A. F. Noor, B. Mehlig, and A. M. Westram. 2017. Interpreting the genomic landscape of speciation: a road map for finding barriers to gene flow. J. Evol. Biol. 30:1450–1477.

Runemark, A., F. Eroukhmanoff, A. Nava-Bolaños, J. S. Hermansen, and J. I. Meier. 2018. Hybridization, sex-specific genomic architecture and local adaptation. Philos. Trans. R. Soc. B Biol. Sci. 373:20170419. The Royal Society.

Sánchez-Guillén, R. A., A. Córdoba-Aguilar, A. Cordero-Rivera, and M. Wellenreuther. 2014. Rapid evolution of prezygotic barriers in non-territorial damselflies. Biol. Linn. Soc. 113:485–496.

Sánchez-Guillén, R. A., A. Córdoba-Aguilar, B. Hansson, J. Ott, and M. Wellenreuther. 2016. Evolutionary consequences of climate-induced range shifts in insects. Biol. Rev. 91:1050–1064.

Sánchez-Guillén, R. A., M. Wellenreuther, J. R. Chávez-Ríos, C. D. Beatty, A. Rivas-Torres, M. Velasquez-Velez, and A. Cordero-Rivera. 2017. Alternative reproductive strategies and the maintenance of female color polymorphism in damselflies. Ecol. Evol.. 7:5592–5602.

Sánchez-Guillén, R. A., M. Wellenreuther, A. Cordero-rivera, and B. Hansson. 2011. Introgression and rapid species turnover in sympatric damselflies. BMC Evol. Biol. 11:210. BioMed Central Ltd.

Sánchez-Guillén, R. A., M. Wullenreuther, and A. Cordero Rivera. 2012a. Strong asymmetry in the relative strengths of prezygotic and postzygotic barriers between two damselfly sister species. Evolution (N. Y). 66:690–707.

Sánchez-Guillén, M. Wellenreuther, and A. Cordero-Rivera. 2012b. Strong asymmetry in the relative strengths of prezygotic and postzygotic barriers between two damselfly sister species. Evolution (N. Y). 66:690–707.

Sweigart, A. L. 2010. Simple Y-autosomal incompatibilities cause hybrid male sterility in reciprocal crosses between *Drosophila virilis* and *D. americana*. Genetics 184:779–787.

Wellenreuther, M., J. Muñoz, J. R. Chávez-Ríos, B. Hansson, A. Cordero-Rivera, and R. A. Sánchez-Guillén. 2018. Molecular and ecological signatures of an expanding hybrid zone. Ecol. Evol. 1–14.

