## Supplemental Information for "Restricted X chromosome introgression and support for Haldane’s rule in hybridizing damselflies"

Table S1. Locations of the sampled sites and respective sample sizes.

| **Latitude** | **Longitude** | **Location** | **Region** | **N** |
| --- | --- | --- | --- | --- |
| 53.24 | -2.58 | Liverpool, UK | allopatric *I. elegans* | 4 |
| 50.53 | 4.43 | Leuven, Belgium | allopatric *I. elegans* | 8 |
| 43.53 | 4.30 | Vigueirat, France | allopatric *I. elegans* | 10 |
| 43.35 | 1.24 | Marais D’Orx, France | allopatric *I. elegans* | 9 |
| 42.38 | 3.01 | Saint Cyprien, France | allopatric *I. elegans* | 10 |
| 42.27 | -8.51 | Cachadas, Spain | allopatric *I. graellsii* | 9 |
| 37.88 | -4.77 | Córdoba, Spain | allopatric *I. graellsii* | 9 |
| 37.10 | 8.33 | Algarve, Portugal | allopatric *I. graellsii* | 12 |
| 37.10 | -8.33 | Riveira de Cobres, Portugal | allopatric *I. graellsii* | 9 |
| 36.28 | 7.22 | Seyhouse, Algeria | allopatric *I. graellsii* | 4 |
| 40.20 | -3.01 | Amposta, Spain | sympatric Central | 10 |
| 39.50 | -0.33 | Carraixet, Spain | sympatric East | 7 |
| 39.07 | -0.31 | Marjal del Moro, Spain | sympatric East | 11 |
| 39.95 | 4.25 | Menorca, Spain | sympatric Menorca | 5 |
| 42.28 | -2.54 | Valpierre, Spain | sympatric North | 10 |
| 42.29 | -2.24 | Las Cañas, Spain | sympatric North | 10 |
| 42.17 | -1.58 | Perdiguero, Spain | sympatric North | 9 |
| 42.24 | -2.42 | Villar, Spain | sympatric North | 9 |
| 42.24 | -2.35 | Valbornedo | sympatric North | 10 |
| 42.48 | -2.58 | Arreo, Spain | sympatric North | 10 |
| 42.30 | -2.52 | Mateo, Spain | sympatric North | 9 |
| 42.37 | -9.23 | Xuño, Spain | sympatric West | 10 |
| 43.29 | -8.19 | Doniños, Spain | sympatric West | 9 |
| 43.62 | -8.11 | Laxe, Spain | sympatric West | 10 |
| 42.69 | -8.66 | Louro, Spain | sympatric West | 10 |
| 36.82 | 11.99 | Pantelleria, Italy | *I. fountaineae* Outgroup | 3 |
| 36.70 | 14.97 | Ispica, Italy | *I. genei* Outgroup | 3 |
| 29.82 | 7.20 | Oued Tissint, Morocco | *I. saharensis* Outgroup | 3 |

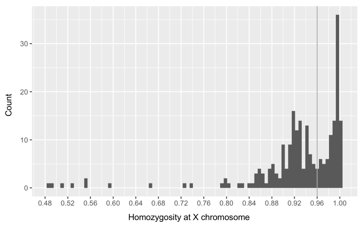

Figure S1. Distribution of observed homozygosity using the full SNP set before filtering of X-linked SNPs. Grey line denotes the chosen cut-off value to classify males and females. The samples at the left tail of the histogram are hybrids (showing intermediate Q values).

Table S2. Statistical results of BGC analysis for the female subset and the complete dataset (females and males).

|  | **females** | | | **females + males** | | |
| --- | --- | --- | --- | --- | --- | --- |
|  | median autosomal | median X-linked | *P*-value | median autosomal | median X-linked | *P*-value |
| Beta | 0.022 | 0.166 | **<0.001** | 0.026 | 0.101 | **<0.001** |
| Alpha | -0.008 | 0.015 | **0.034** | -0.017 | 0.015 | **<0.001** |

Table S3. Statistical results of ABBA-BABA analysis for the female subset and the complete dataset (females and males).

|  | *I. elegans* | | | | | |
| --- | --- | --- | --- | --- | --- | --- |
|  | **females** | | | **females + males** | | |
| Included individuals | median autosomal | median X-linked | *P*-value | median autosomal | median X-linked | *P*-value |
| Q = 0 | 0.005 | -0.009 | **<0.001** | 0.003 | -0.006 | **<0.001** |
| Q < 0.1 | 0.018 | 0.002 | **<0.001** | 0.014 | -0.001 | **<0.001** |
| Q < 0.25 | 0.026 | 0.009 | **<0.001** | 0.018 | 0.004 | **<0.001** |
|  | *I. graellsii* | | | | | |
|  | **females** | | | **females + males** | | |
| Included individuals | median autosomal | median X-linked | *P*-value | median autosomal | median X-linked | *P*-value |
| Q = 0 | -0.003 | -0.008 | **0.007** | -0.002 | -0.007 | **<0.001** |
| Q > 0.9 | 0.005 | -0.007 | **<0.001** | 0.005 | -0.005 | **<0.001** |
| Q > 0.75 | 0.009 | 0.001 | **<0.001** | 0.009 | 0.001 | **<0.001** |
